# Engineering Microbial R- and S- β-Hydroxybutyrate Production

**DOI:** 10.1101/2025.07.15.664879

**Authors:** Burak Kizil, Sinem Ulusan, Aydan Torun, Mehmet Gumustas, Şeniz Yuksel, Sreeparna Banerjee

## Abstract

Beta-hydroxybutyrate (BHOB) is a therapeutically valuable enantiomeric ketone body that is synthesized by both prokaryotes and eukaryotes. Bacterial synthesis of BHOB has so far mostly been exploited for the synthesis of biofuels and to a lesser extent for the pharmaceutical industry. In this study, we carefully evaluated the expression, induction, selection, and detection elements and identified multigene pathways to direct the synthesis of physiologically relevant amounts of BHOB. Utilizing the reversal of the beta-oxidation pathway that has previously been used for biofuel generation, we engineered *E. coli* strains with a prebiotic-activated circuit to secrete either (*S*)-BHOB or (*R*)-BHOB. Our findings demonstrate high-yield microbial BHOB production with a defined enantiomeric composition and highlight its therapeutic potential for cancer and gut–brain axis involving disorders.

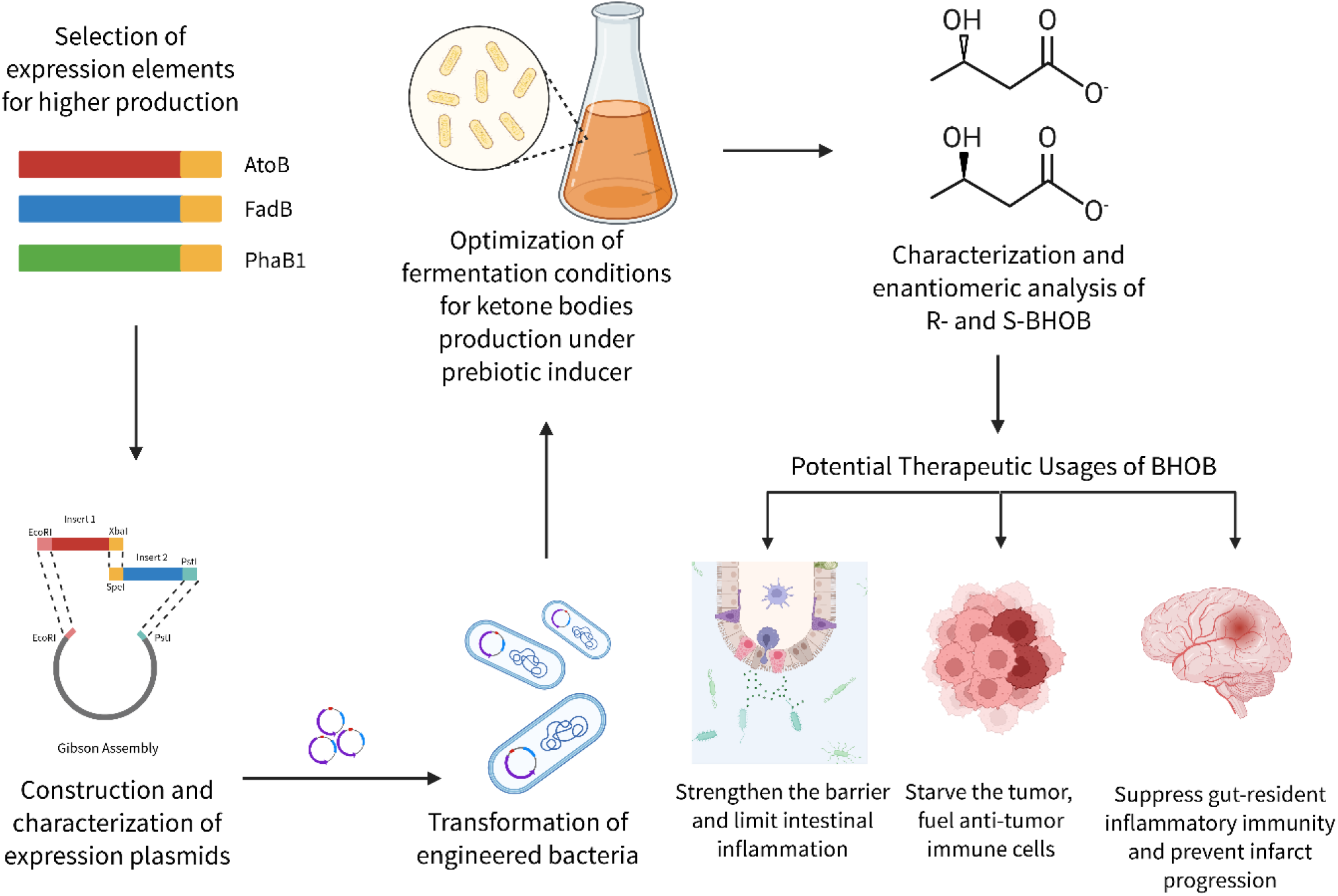

## Introduction

Advancement in synthetic biology has allowed the engineering of commensal microorganisms into probiotics that can restore homeostasis within the gastrointestinal tract^1^. Such Next-Generation Probiotics (NGP) have gained popularity for the treatment of diseases such as infections, metabolic disorders, inflammation, and cancer^2^. For example, *Saccharomyces boulardii* has been engineered to treat metabolic disorders^3^ while *Escherichia Coli Nissle* 1917 has been engineered to ameliorate phenylketonuria (PKU), bacterial infections, and cancer^4^.

In humans, ketone bodies are synthesized from acetyl CoA that is derived from β-oxidation of fatty acids.^5^ Among them, Acetoacetate (AcAc) and β-hydroxybutyrate (BHOB) are the primary ketone bodies that can be metabolized for energy generation and have signaling functions. The primary organ for BHOB synthesis is the liver and 3-hydroxymethylglutaryl-CoA synthase (HMGCS2), the rate-limiting enzyme for its synthesis, is highly expressed in hepatocytes.^6^ Ketone bodies are transported from the liver via the circulation to extra hepatic tissues where they can be converted to acetyl CoA by 3-oxoacid CoA-transferase 1 (OXCT1) for metabolism via the TCA cycle.^6^ When glucose availability is limited, neurons, skeletal muscles and renal epithelial cells rely on ketone bodies as an alternative source of ATP.

Humans can only metabolize (*R*)-BHOB to acetyl CoA and ATP; moreover, any available (*S*)-BHOB is known to be converted to (*R*)-BHOB^7^. The binding of BHOB to its receptors, however, is non-stereoselective, suggesting that both (*S*)-and (*R*)-enantiomers can mediate the signaling functions of BHOB^7^. Microorganisms are known to synthesize and secrete BHOB via the direct condensation of two acetyl coenzyme A (CoA) molecules with subsequent reduction steps^8^.

In the current study, we describe a strategy to produce physiologically relevant amounts of *(S*)-and (*R*)-β-hydroxybutyrate (BHOB) via the reversal of the β-oxidation pathway in *E. coli* K12. It is known that wild type *E. coli* can utilize carbon sources (glucose, glycerol, etc.) and produce ethanol, acetate and lactate under anaerobic conditions^9^. To direct the carbon atoms towards the synthesis of BHOB, we used the *E. coli* strain K12 MG1655 that was modified by eliminating the enzymes required for the production of lactate, ethanol, fumarate, and various aldehyde pathways (ΔldhA, ΔadhE, ΔackA, Δpta, ΔfrdC)^10^. This ensured that carbon atoms from glycerol would be directed towards ketone body synthesis rather than glycolysis. This strategy has been extensively used in studies relevant to bio-fuel production^11^. In addition, *E. coli* strains form a considerable proportion of the gut microbiota^12^. Therefore, in this study carried out genetic manipulation of *E. coli* K12 commensal strain for large-scale metabolite production^13^.

Microbial synthetic biology enables the engineering of probiotic bacteria to produce beneficial metabolites *in situ^14^*. A circuit for BHOB expression was constructed using the most effective pre-characterized expression elements selected from the International Genetically Modified Organisms (iGEM) library. To generate (*S*)-β-hydroxybutyrate, *atoB* and *fadB* were selected. To generate (*R*)-β-hydroxybutyrate, *atoB* and *phaB* were overexpressed. The (*R*)-β-hydroxybutyryl CoA or (*S*)-β-hydroxybutyryl CoA can be converted to the respective β-hydroxybutyrate via the enzymatic action of several endogenous thioesterases expressed in *E. coli* ^10,11,15^.

Our goals were to maximize bacterial BHOB synthesis and secretion, characterize enantiomeric purity, and discuss the therapeutic implications of gut-derived BHOB. We demonstrate that engineered *E. coli* K12 can secrete millimolar amounts of BHOB, and we discuss how such microbial BHOB might impact diseases such as cancer and stroke by modulating immune and metabolic pathways.

## Materials and Methods

### Rational design of the enzyme circuit

AtoB and FadB were selected for the synthesis of (*S*)-BHOB (Figure 1). AtoB is a thiolase that can catalyze the condensation of acetyl-CoA with acyl-CoA of various chain lengths but has higher specificity for short-chain acyl-CoA molecules^15^. FadB, also known as β-Hydroxyacyl-CoA dehydrogenase, is a reversible enzyme that possesses both hydroxyacyl-CoA dehydrogenase and enoyl-CoA hydratase activity with broad chain-length specificity. The enzyme uses NADH to generate acetyl-CoA in each cycle of β-oxidation. FadB is known to strictly generate the (*S*)-stereospecific isomer of 3-hydroxybutyryl-CoA^15^. AtoB and PhaB1 were selected for the synthesis of (*R*)-BHOB. PhaB1 [3-oxacyl (acyl carrier protein) reductase 2] is an enantioselective β-Hydroxyacyl-CoA dehydrogenase, producing the (*R*)-enantiomer in higher quantities than the (*S*)-enantiomer ^16^. PhaB [3-oxacyl (acyl carrier protein) reductase 2, E.C. 1.1.1.100] converts acetoacetyl CoA to *R*-β-hydroxybutyryl CoA. This enzyme is well-described and characterized for the generation of (*R*)-β-hydroxybutyrate in the iGEM library.

**Figure 1.**
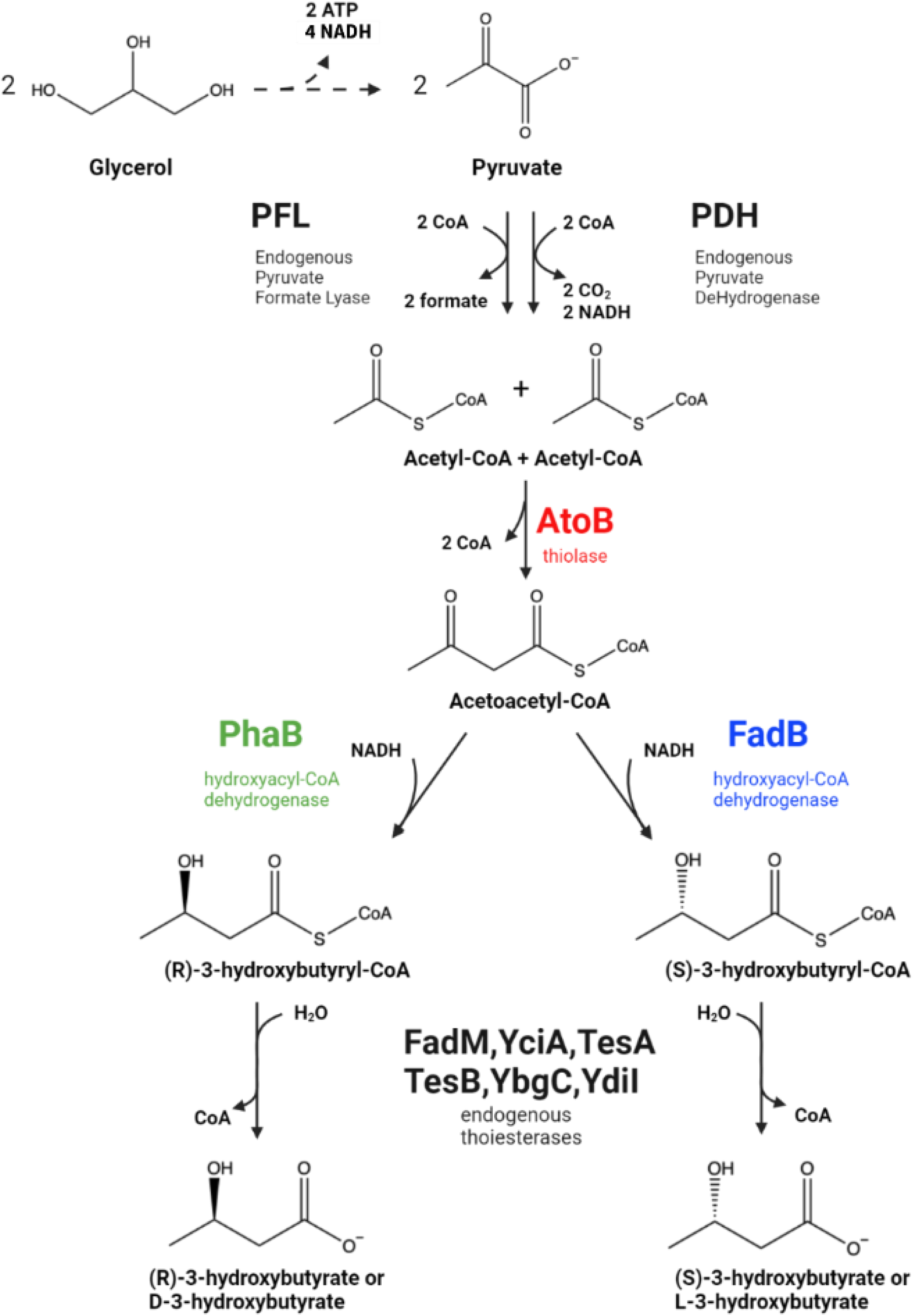
Schematic representation of the synthesis of (*S*)-BHOB and (*R*)-BHOB from glycerol in *E. coli*. PhaB corresponds to (R)-3HB-CoA dehydrogenase from *R. eutropha* H16 (isolated from the construct described AtoB, Thiolase from *E. coli* MG1655; FadB, 3-Hydroxacyl-CoA Dehydrogenase from *E. coli* MG1655.

### Plasmid construction

Enhancement of expression was ensured with the inclusion of a strong ribosome binding site (RBS) with a translation rate of 0.2059 min^-1^ (http://parts.igem.org/Part:BBa_B0034) and strong pBAD promoter (responsive to low concentrations of prebiotic called *L*-arabinose, http://parts.igem.org/Part:BBa_K206000) upstream of the enzymes. Each enzyme was tagged with 10x Histag (http://parts.igem.org/Part:BBa_K844000) for detection by western blot. In vivo assembly and characterization of the functional pathway were done on pUC19 derived high copy number cloning and expression vectors (pSB1C3).

All enzymes were cloned into the plasmid pSB1C3, a pUC19 derived high copy number cloning vector that is designed for consecutive cloning. The pSB1C3 vector consists of a multiple cloning site bearing restriction enzyme sites for *EcoRI, XbaI, SpeI* and *PstI*. C-terminal cloning in this vector was carried out by using the restriction sites *SpeI* and *PstI* in the vector and *XbaI* and *PstI* sites in the insert. Using the Gibson cloning technique, an insert bearing *XbaI* and *PstI* sticky ends was ligated to the *SpeI* and *PstI* sites, respectively (Figure 2A). When the *XbaI*-*SpeI* sites were ligated, a ‘TACTAG’ sequence was generated, which could not be digested with the same enzymes again. The *XbaI* site, downstream of *EcoRI* in the vector and the *SpeI* site upstream of *PstI* on the insert were preserved and could be used for future cloning experiments. The sequences necessary for the overexpression of the selected genes and their detection after overexpression were identified from an iGEM catalog. Sequences of the arabinose responsive pBAD promoter (http://parts.igem.org/Part:BBa_K206000), ribosome binding side (RBS) (http://parts.igem.org/Part:BBa_B0030), and a 10x Histidine tag with a terminator sequence (http://parts.igem.org/Part:BBa_K844000) were used as template or extensions at the flanking ends of primers. Sequences of *atoB, phaB1* (AM260479.1; *Ralstonia eutropha* H16 chromosome 1; http://parts.igem.org/Part:BBa_K934001) and *fadB*, (GenBank: U00096.2, NCBI) from the *E. coli* strain K12 MG1655, were amplified from the genomic DNA by nested PCR. After each cloning, the products were transformed and amplified in the *E. coli* DH5alpha strain and confirmed by PCR and Sanger sequencing.

**Figure 2.**
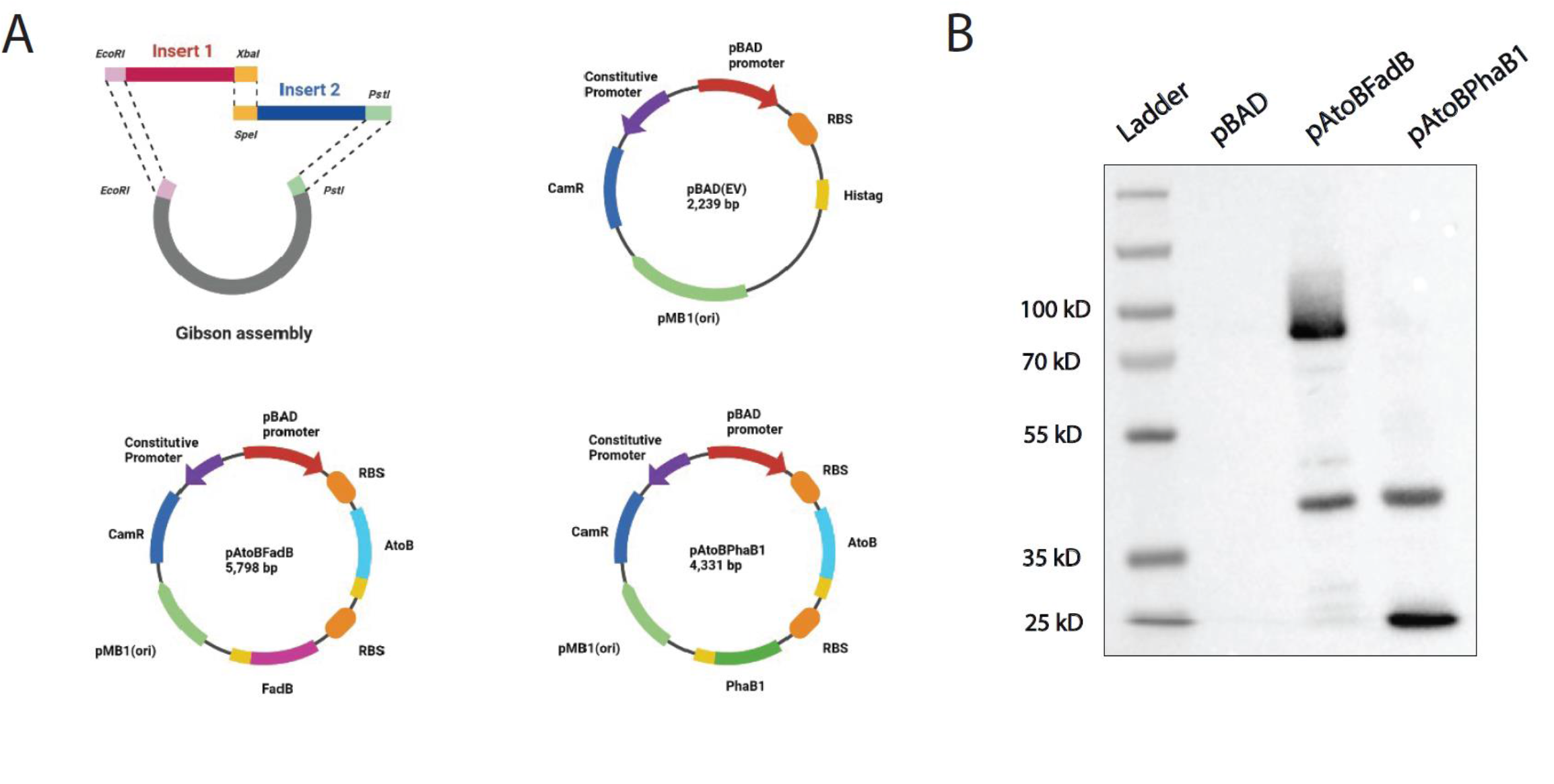
Cloning strategy and expression of genes for the synthesis of (*R*)-BHOB and (*S*)-BHOB. **A.** Construction of the vector with an inducible pBAD promoter, ribosome binding site upstream of each gene of interest and a Histidine tag (Histag, yellow) downstream of each gene of interest. The pAtoBFadB vector was designed for the synthesis of (*S*)-BHOB while pAtoBPhaB1 was designed for the synthesis of (*R*)-BHOB. **B**. Western blot with an anti-histidine antibody showing the expression of FadB (72 kD) and AtoB (42 kD) in bacteria transformed with the pAtoBFadB vector and the expression of AtoB and PhaB1 (25 kD) in bacteria transformed with the pAtoBPhaB1 vector.

### Bacterial culture protocol

The bacterial strains used in the current study are shown in Table 2. Five mL of pre-cultures for the bacterial strains pBAD, pAtoBFadB and pAtoBpHaB1 were prepared using Luria Broth (LB) supplemented with Chloramphenicol at a final concentration of 25 µg/mL. The cultures were grown overnight at 37 °C with shaking at 180 rpm. To scale up, 500 µL of pre-culture was inoculated into 50 mL of LB supplemented with chloramphenicol at a final concentration of 25 µg/mL. The inoculated cultures were grown in pre-autoclaved 250 mL glass Erlenmeyer flasks at 37°C and 180 rpm until an OD_600_ of 0.5-0.6 was reached. The culture was next induced with 50 µL of L-Arabinose from a stock solution 10% (w/v). After 4 hours of induction, the cultures were centrifuged at 5000 x *g* for 10 min, and the supernatants were discarded. The pellets were carefully resuspended in 20 mL of Terrific Broth (TB) to prepare for fermentation. Meanwhile, 25 mL glass Erlenmeyer flasks containing 50 mg CaCO_3_ were autoclaved and dried at room temperature. The resuspended pellets (20 mL) were transferred to the 25 mL Erlenmeyer flasks and supplemented with 5 mM calcium pantothenate, 2 mM MgSO_4_, 5 mM (NH_4_)_2_SO_4_, 100 µM FeSO_4_ and 20 µL L-Arabinose (10% w/v), in addition to 25 µg/mL of chloramphenicol. The cultures were incubated at 37°C with shaking at 180 rpm before closing the lids tightly so that air exchange was prevented. After 36-48 hours of incubation the cultures were collected and centrifuged at 5000 x *g* for 30 min. The supernatants were collected, and filter sterilized with 0.44 μm filters for further analyses. The weight of the pellets was used to determine the production efficiency of BHOB.

### Protein isolation from bacteria

To confirm the expression of AtoB, FadB and PhaB, total protein was isolated from the respective bacterial strains. For this, aliquots from overnight cultures (1 mL) were centrifuged at 14000 rpm for 1 min. The supernatant was discarded, and the pellet was resuspended in TEG buffer (25 mM Tris-Cl, 50 mM Glucose, 10 mM EDTA, pH adjusted to 8.0) at a final concentration of 100 mg pellet/mL. The suspensions were mixed until the clumps had disappeared. TEG lysis buffer containing lysozyme (0.5 mg/ml final concentration) was added as 10% (v/v) of the suspension, mixed thoroughly and incubated at 37°C for 30 minutes. Equal volume of each sample was mixed with sample loading dye [1.1 mL 4 X running buffer (90.85 gr. Tris-Base, 20 mL 10% SDS, dH_2_O to 500 mL, pH adjusted to 8.8), 1.75 mL Glycerol, 0.5 mL 2-mercaptoethanol, 0.25 mL 0.2% Bromophenol blue, 20% SDS and dH_2_O up to 10 mL], incubated at 100°C for 10 min. The samples were cooled and loaded on a 10% SDS-PAGE gel for western blot using an anti-histidine tag antibody.

### Gas Chromatography-Mass Spectrometry (GC-MS) for evaluation of ketone bodies

#### Derivatization and Sample Preparation for GC-MS

The conditioned media collected from the culture of pBAD, pAtoBFadB and pAtoBpHaB1 strains (Table 2) were derivatized to solubilize the water-soluble components (including ketone bodies and acetate) in hexane^11^. Methyl chloroformate (MCF) was used to obtain more stable propyl derivatives rapidly under fume hood. For this, 300 μL of conditioned medium was taken into 2 mL eppendorf tubes and mixed with 5 μg/mL heptanoic acid as the internal standard. Propanol, pyridine and MCF (3:2:1, v/v/v, 500 μL) was mixed with the conditioned medium and ultrasonicated for 1 min. The derivatized components were extracted by the addition of 500 μL gas chromatography (GC) grade hexane and vortexed for 1 minute. The tube was centrifuged at 2000 x *g* for 5 min and the organic phase (upper layer) was collected in a fresh glass tube containing 10 mg anhydrous sodium sulfate (desiccant). A second aliquot of 500 μL of hexane was mixed with the derivatized solution, centrifuged at 2000 x *g* for 5 min and the organic phase was again collected for GC. To quantify the BHOB, a calibration curve was generated using the sodium salt of 3-hydroxybutyrate and acetate. Aqueous solutions of these salts were prepared at a final concentration of 0.1 mM, 1 mM and 10 mM in water and derivatized as above to generate the calibration curve.

#### Sample Analysis with GC-MS

GC-MS was carried out with a ZB-FFAP column (length, 30 m; diameter, 0.32 mm; film, 0.50 µm; Phonemenex Inc., USA) to detect and quantify BHOB and acetate in the bacterial conditioned medium. The injection port temperature was set at 150 °C. Helium was used as a carrier gas and the oven temperature was held at 80 °C for 1 min. The temperature was then elevated to 135 °C at a rate of 10 °C/min and kept at that temperature for 5 min. The final temperature was adjusted to 230 °C by increasing the temperature at a rate of 30 °C/min and held for 6 min. Ionization temperature was set at 230 °C for mass spectrometry. 4 µL of sample was injected in a splitless mode. The GC-MS experiments were carried out at UNAM, Bilkent University.

### Chiral High Performance Liquid Chromatography (HPLC)

#### Methyl esterification of 3-hydroxybutyrate

50 mL of the conditioned media from pBAD, pAtoBFadB and pAtoBpHaB1 strains (Table 2) were filter sterilized and heated at 50 °C for evaporation on a heat block. The resulting dark brown residue was resuspended in 2 mL acidic methanol (methanol:concentrated HCl at a ratio of 4:1 v/v). The acidic methanol solution was then transferred to a sealed tube and heated at 100 °C for 3 hours in a water bath. After cooling to room temperature, 200 µL of this solution was mixed with 1 mL of isopropanol and vortexed. The mixture was then centrifuged at 13,000 x *g* for 10 min and the precipitate was removed. The supernatant containing methyl esterified BHOB was used for the chiral HPLC analysis^11^. Commercially obtained methyl esterified (R)-BHOB (Sigma, Cat No: 298360) and (S)-BHOB (Cayman, Cat No: 21822) were used to generate the calibration curves.

#### Sample Analysis with chiral HPLC

Chiral HPLC analysis was carried out with a Lux Cellulose 1 – 250 × 4.6 mm (3 µm particle size) column (Phenomenex, Torrance, USA). Column temperature was kept at 40 °C and mobile phase composition was n-hexane/isopropanol (9:1) at a flow rate of 0.7 mL/min. 5 μL samples were injected to the column under 45 bar pressure. The UV detector was set at 210 nm for the detection of methyl esterified BHOB. Commercially sources methyl esters of (*R*)-BHOB and (*S*)-BHOB were used as standards with retention times of 8.15 and 10.31 min respectively.

## RESULTS

### Selection of elements of the expression vector, cloning and gene expression

Effective enzyme production requires a strong ribosome binding site (RBS) on the plasmid. For the synthesis of *atoB, fadB* and *phaB*, a strong RBS characterized with a translation rate of 0.2059 min^-1^ was selected from the IGEM repository and cloned upstream of each gene of interest (Figure 2A).

No secondary structure was expected in the given ribosome binding region. The promoter selected for the construct was pBAD, a well-characterized *E. coli* promoter that can be induced in the presence of L-arabinose^17^. This promoter was specifically designed to be responsive to low concentrations of L-arabinose but is known to be highly repressed in the presence of glucose^17^. We also observed very low expression of genes cloned downstream of the pBAD promoter when glucose was used in the bacterial culture medium (data not shown). Therefore, glycerol was used as the carbon source for all experiments.

The Gibson cloning method was used to clone histidine tagged *atoB* and *fadB* or *atoB* and *phaB* downstream of their individual RBS and under the control of the inducible promoter pBAD. The plasmid was then transformed into the *E. coli* strain BE1579 (*E. coli* K12 MG1655substrain) (Table 2). The bacterial lysate was collected and evaluated for the expression of AtoB, FadB and PhaB by western blot using an anti-histidine antibody. The presence of a band at the expected size of the protein product was considered as confirmation of expression (Figure 2B).

### Characterization of the secreted BHOB

The amount of BHOB secreted into the conditioned medium collected from the strains pBAD (Empty vector, EV), pAtoBFadB [secreting (*S*)-BHOB] and pAtoBpHaB1 [secreting (*R*)-BHOB] (please see Table 2 for a description of these strains) was next determined using Gas Chromatography-Mass Spectrometry (GC-MS) (Table 1).

**Table 1.** Amount of BHOB and acetic acid secreted into the conditioned medium of bacterial strains.

|  | BHOB (mM) | Acetic Acid (mM) |
| --- | --- | --- |
| pBAD (EV) | $0.04 \pm 0.007$ | $4.68 \pm 0.007$ |
| pAtoBpHaB | $31 \pm 14.329$ | $4.57 \pm 1.039$ |
| pAtoBFadB | $26 \pm 13.650$ | $3.49 \pm 0.849$ |

**Table 2.** Bacterial strains used in this study.

| Strains | Description | Source |
| --- | --- | --- |
| DH5α | fhuA2 $\Delta$(argF-lacZ) U169 phoA glnV44 Φ80Δ (lacZ)M15 gyrA96 recA1 relA1 endA1 thi-1 hsdR17 | Laboratory Collection |
| K12<br>MG1655 | F- lambda- ilvG- rfb-50 rph-1 | Laboratory Collection |
| BE1579 | *E. coli* MG1655 DadhE : : FRT, DldhA : : FRT, DackA-pta : : FRT, DfrdC : : FRT | Thomas Bobik <sup>10</sup> |
| pBAD | *E. coli* MG1655 DadhE : : FRT, DldhA : : FRT, DackA-pta : : FRT, DfrdC : : | This study |
| pAtoBFadB | *E. coli* MG1655, DadhE : : FRT, DldhA : : FRT, DackA-pta : : FRT, DfrdC : : | This study |
| pAtoBpHaB1 | *E. coli* MG1655, DadhE : : FRT, DldhA : : FRT, DackA-pta : : FRT, DfrdC : : | This study |

The chromatograms showing the synthesis and secretion of BHOB from *E. coli* strain BE1579 transformed with pAtoBFadB or pAtoBpHaB are shown in Figure 3. No synthesis of BHOB was observed in the strain transformed with the empty vector (EV).

**Figure 3.**
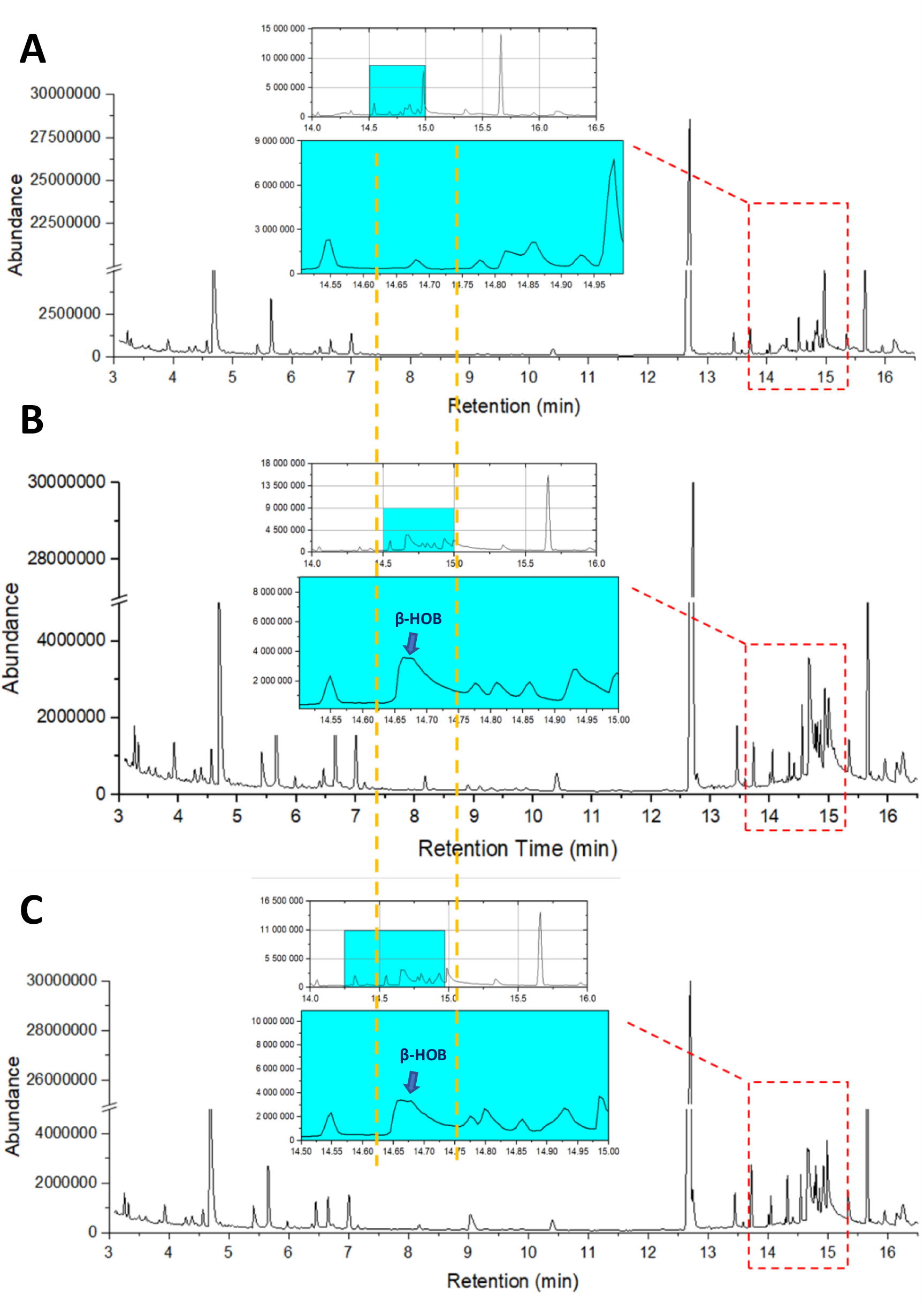
GC-MS showing the secretion of beta-hydroxybutyrate from *E. coli* BE1579 bacteria. BHOB had a retention time of 14.628 min **A.** The bacteria containing the empty vector (pBAD) secreted a negligible amount of BHOB. **B & C.** The bacteria transformed with pAtoBFadB (**B**) and pAtoBpHaB1 (**C**) were found to secrete between 10 mM – 45 mM BHOB.

To determine the enantiomeric ratio [(*R*)-BHOB: (*S*)-BHOB)] of the secreted BHOB, chiral HPLC analysis was carried out. The samples were methyl esterified prior to loading as carboxylic acids tend to retain in the column and are harder to separate. The chiral HPLC chromatograms are shown in Figure 4. The enantiomeric excess of either (*R*)-BHOB and (*S*)-BHOB was calculated from the area under the corresponding peaks. The area under the curve obtained from chiral HPLC was applied to the concentrations obtained from the corresponding GC-MS runs to calculate the amounts of (*R*)-BHOB and (*S*)-BHOB secreted in mM. The strain with pAtoBpHaB1 was found to secrete mostly (*R*)-BHOB at an *R*:*S* ratio of 14:1 and while the strain with pAtoBFadB was found to express mostly (*S*)-BHOB at an *R*:*S* ratio of 1:63.

**Figure 4.**
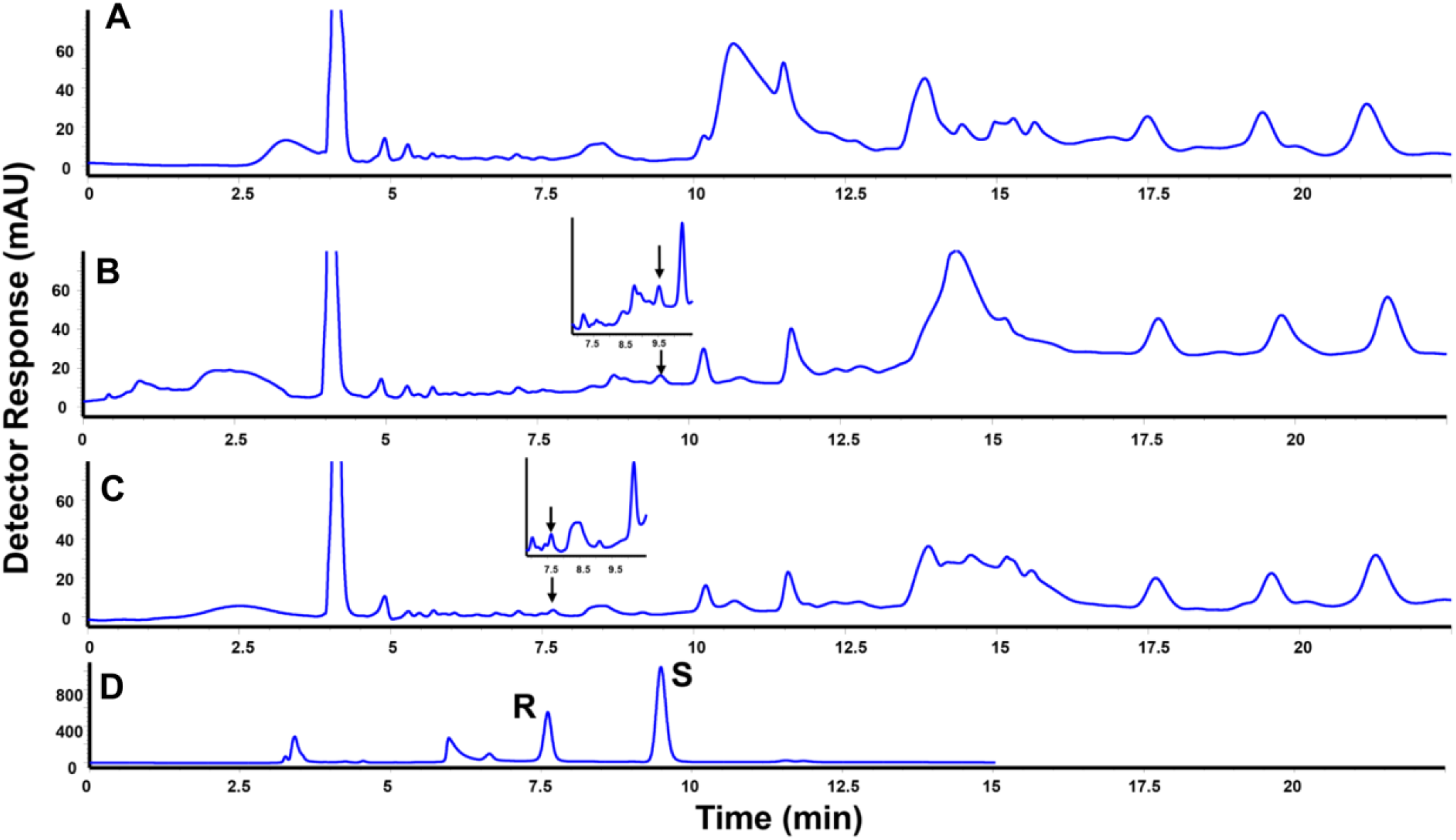
Secretion of (*R*)-BHOB and (*S*)-BHOB from the engineered *E. coli* strain. Chiral HPLC chromatograms of the conditioned medium from bacteria transformed with **A**. pBAD, **B**. pAtoBpHaB1 showing a peak for (*R*)-BHOB in the inset and **C**. pAtoBFadB showing a peak for (*S*)-BHOB in the inset. **D**. Standard (*R*)-BHOB and (*S*)-BHOB peaks generated with a racemic mixture of the pure compounds.

## Statistical analyses

All experiments were repeated at least 3 times independently, unless otherwise stated. The data were compared with either Students t-test or ANOVA, followed by Tukey’s post hoc test. Data analysis and graphing was carried out with Graph Pad Prism (La Jolla, CA, USA). P <0.05 was statistically significant.

## Discussion

We have demonstrated that *E. coli* can be engineered to secrete high levels of either (R)-or (S)-β-hydroxybutyrate. The synthesized BHOB concentrations (∼30 mM) are comparable to physiologically relevant ketone levels, indicating the utility of this approach for probiotic delivery of ketones *in vivo*. Below we discuss potential implications of bacterially derived BHOB for host epithelial, immune, and neural health, especially in the context of stroke and cancer. Future studies on engineering probiotic bacteria that can secrete BHOB, colonization of such bacteria in animal models and evaluation of signaling pathways activated will be necessary to fully evaluate the benefits of BHOB secreting bacteria on human health.

### BHOB and Epithelial/Immune Homeostasis

BHOB has emerging roles in modulating immune and epithelial physiology^18^. In models of intestinal inflammation, BHOB promotes tissue repair. For example, in murine colitis, rectal BHOB treatment was shown to drive macrophages toward an anti-inflammatory (M2) phenotype via STAT6 activation, which in turn enhanced intestinal epithelial regeneration^18^. Thus, gut-derived BHOB may strengthen the epithelial barrier and limit intestinal inflammation by skewing macrophages toward a tissue-repairing state^18^. These findings suggest that an engineered probiotic that secretes BHOB could help maintain gut integrity and immune balance in inflammatory bowel diseases.

### Stroke

Following ischemic stroke, BHOB levels were shown to rise significantly in both the blood and brain^19^. In experimental models such as middle cerebral artery occlusion (MCAO), BHOB increases in the liver, circulation, and ischemic brain regions, while glucose levels simultaneously decline^19^. Similarly, in stroke patients, elevated serum and urinary BHOB levels were observed within 72 hours post-stroke, returning to baseline by the chronic phase (3–6 months)^19^. Exogenous BHOB administration was shown to be neuroprotective by reducing infarct size and improving neurological function^19^. Mechanistically, BHOB exerts its effects through multiple pathways: stabilizing mitochondrial function, reducing oxidative stress, inhibiting NLRP3 inflammasome activation, protecting the blood–brain barrier, and regulating gene expression via histone deacetylase (HDAC) inhibition^19^.

Beyond exogenous supplementation, BHOB can also be elevated through metabolic interventions^19^. Experimental induction of ketosis—via short-term fasting combined with a ketogenic diet (KD) or subacute KD feeding—has demonstrated beneficial effects in stroke models. These include enhanced motor recovery, reduced infarct volume, improved cerebral blood flow, and protection against mitochondrial damage^19^. Interestingly, recent studies have linked BHOB to the regulation of γδ T cells, which play a critical role in post-stroke inflammation^20^. Shichita et al. (2009) showed that γδ T cells are key contributors during the subacute phase of stroke, and their depletion can lead to a ∼50% reduction in infarct size^21^. Further research by Benakis et al. revealed that these cells originate from the gut and migrate to the brain following ischemic injury, where they amplify neuroinflammation^22^. The ketogenic diet—by elevating BHOB—modulates γδ T cell activity in a time-and context-dependent manner^19^. While short-term or moderate KD promotes expansion of metabolically protective γδ T cells in visceral adipose tissue, prolonged ad libitum KD leads to metabolic dysfunction and depletion of these cells^20^. Given the pathogenic role of γδ T cells in stroke, sustained BHOB exposure may paradoxically limit their pro-inflammatory effects by depleting their pool, thus contributing to neuroprotection. Building on this concept, our engineered BHOB-producing bacteria represent a novel microbiome-based therapeutic strategy. By delivering BHOB timely and locally within the gut, these microbes have the potential to modulate γδ T cell activity, and limit neuroinflammation and infarct expansion after stroke.

### Cancer

Tumor cells often depend on glucose, creating an immunosuppressive microenvironment rich in lactate^23^. Ketone bodies are generally poor fuels for many cancers, making high BHOB/low glucose conditions (as in ketogenic diets) selectively stressful to tumors^23^. In murine cancer models, ketogenic regimens decrease the levels of lactate and inhibit cancer cell proliferation while enhancing cytotoxic immune responses^23^. For example, combining ketone supplementation with checkpoint blockade was shown to synergistically improve survival in multiple tumor models, even in some therapy-resistant tumors^23^.

At the cellular level, BHOB can influence immune checkpoints and metabolism^23^. BHOB inhibits histone deacetylases and promotes the expression of genes involved in memory T-cell formation^23^. In the tumor microenvironment, BHOB was reported to increase expression of antigen-presenting machinery and boost CD8^+^ T-cell activity, while reducing suppressive cell types^23^. Moreover, KD-induced BHOB was shown to lower intratumoral lactate, relieving lactate-mediated T-cell suppression^23^. Locally produced BHOB could starve tumors and concurrently “fuel” anti-tumor immune cells. Overall, engineered probiotic BHOB production might serve as a cancer metabolic therapy by depriving cancer cells of glucose, altering the acidic microenvironment, and potentiating cytotoxic immune infiltrates.

### Future Therapeutic Perspectives

Our engineered *E. coli* strain provides a platform for continuous gut-derived ketone delivery. Unlike oral ketone esters, a living probiotic could sustain moderate BHOB levels with minimal dosing. Before clinical translation, in vivo studies will be needed to confirm colonization, BHOB production in the gut, and systemic uptake. It will also be important to examine effects on gut microbiota and any off-target metabolites. Nevertheless, the high yield and enantiomeric control achieved here are promising. Given the diverse roles of BHOB in immunity, neuroprotection, and metabolism, microbial BHOB production could be harnessed for novel therapies in stroke and cancer.

## Acknowledgements

The authors would like to acknowledge Drs. Haluk Hamamci, Salih Ozcubukcu, Huseyin Avni Oktem, and Gorkem E. Gunbas (Middle East Technical University, Ankara, Turkiye) and Sibel Özkan (Faculty of Pharmacy, Ankara University) for sharing resources. Dr. Omer H. Yilmaz (Koch Institute for Integrative Cancer Research at MIT) is gratefully acknowledged for discussions and feedback. Dr. Thomas A. Bobik (Iowa State University, Ames, IA) is gratefully acknowledged for the generous donation of the BE1579 *E. coli* strain. We also acknowledge the technical assistance provided by Rezwan Siddiqui, Erdem Ercan and Rajeshwari Choudhury Nath. Zeynep Erdogan at UNAM, Bilkent University, is gratefully acknowledged for her help with the GC-MS analyses. This project was supported by TUBITAK (118Z170) and AdımODTU.

## References

(1) Aggarwal, N.; Breedon, A. M. E.; Davis, C. M.; Hwang, I. Y.; Chang, M. W. Engineering Probiotics for Therapeutic Applications: Recent Examples and Translational Outlook. Curr Opin Biotechnol 2020, 65, 171–179. 10.1016/j.copbio.2020.02.016.

(2) Langella, P.; Guarner, F.; Martín, R. Editorial: Next-Generation Probiotics: From Commensal Bacteria to Novel Drugs and Food Supplements. Front Microbiol 2019, 10, 1973. 10.3389/fmicb.2019.01973.

(3) Durmusoglu, D.; Al’Abri, I. S.; Collins, S. P.; Cheng, J.; Eroglu, A.; Beisel, C. L.; Crook, N. In Situ Biomanufacturing of Small Molecules in the Mammalian Gut by Probiotic Saccharomyces Boulardii. ACS Synth Biol 2021, 10 (5), 1039–1052. 10.1021/acssynbio.0c00562.

(4) Crook, N.; Ferreiro, A.; Gasparrini, A. J.; Pesesky, M. W.; Gibson, M. K.; Wang, B.; Sun, X.; Condiotte, Z.; Dobrowolski, S.; Peterson, D.; Dantas, G. Adaptive Strategies of the Candidate Probiotic E. Coli Nissle in the Mammalian Gut. Cell Host Microbe 2019, 25 (4), 499-512.e8. 10.1016/j.chom.2019.02.005.

(5) Puchalska, P.; Crawford, P. A. Metabolic and Signaling Roles of Ketone Bodies in Health and Disease. Annu Rev Nutr 2021, 41 (1), 49–77. 10.1146/annurev-nutr-111120-111518.

(6) Berg, J. M.; Tymoczko, J. L.; Stryer, L. Biochemistry, 12th Editi.; Freeman: New York, 2012.

(7) Newman, J. C.; Verdin, E. β-Hydroxybutyrate: A Signaling Metabolite. Annu Rev Nutr 2017, 37, 51–76. 10.1146/annurev-nutr-071816-064916.

(8) Kocharin, K.; Chen, Y.; Siewers, V.; Nielsen, J. Engineering of Acetyl-CoA Metabolism for the Improved Production of Polyhydroxybutyrate in Saccharomyces Cerevisiae. AMB Express 2012, 2 (1), 52. 10.1186/2191-0855-2-52.

(9) Martínez-Gómez, K.; Flores, N.; Castañeda, H. M.; Martínez-Batallar, G.; Hernández-Chávez, G.; Ramírez, O. T.; Gosset, G.; Encarnación, S.; Bolivar, F. New Insights into Escherichia Coli Metabolism: Carbon Scavenging, Acetate Metabolism and Carbon Recycling Responses during Growth on Glycerol. Microb Cell Fact 2012, 11 (1), 46. 10.1186/1475-2859-11-46.

(10) Volker, A. R.; Gogerty, D. S.; Bartholomay, C.; Hennen-Bierwagen, T.; Zhu, H.; Bobik, T. A. Fermentative Production of Short-Chain Fatty Acids in Escherichia Coli. Microbiology (United Kingdom) 2014, 160 (PART 7), 1513–1522. 10.1099/mic.0.078329-0.

(11) Tseng, H.-C.; Martin, C. H.; Nielsen, D. R.; Prather, K. L. J. Metabolic Engineering of Escherichia Coli for Enhanced Production of (R)-and (S)-3-Hydroxybutyrate. Appl Environ Microbiol 2009, 75 (10), 3137–3145. 10.1128/AEM.02667-08.

(12) Tenaillon, O.; Skurnik, D.; Picard, B.; Denamur, E. The Population Genetics of Commensal Escherichia Coli. Nat Rev Microbiol 2010, 8 (3), 207–217. 10.1038/nrmicro2298.

(13) Marisch, K.; Bayer, K.; Cserjan-Puschmann, M.; Luchner, M.; Striedner, G. Evaluation of Three Industrial Escherichia Coli Strains in Fed-Batch Cultivations during High-Level SOD Protein Production. Microb Cell Fact 2013, 12 (1), 58. 10.1186/1475-2859-12-58.

(14) Yadav, M.; Shukla, P. Efficient Engineered Probiotics Using Synthetic Biology Approaches: A Review. Biotechnol Appl Biochem 2020, 67 (1), 22–29. 10.1002/bab.1822.

(15) Clomburg, J. M.; Vick, J. E.; Blankschien, M. D.; Rodríguez-Moyá, M.; Gonzalez, R. A Synthetic Biology Approach to Engineer a Functional Reversal of the β-Oxidation Cycle. ACS Synth Biol 2012, 1 (11), 541–554. 10.1021/sb3000782.

(16) Silvestrini, L.; Drosg, B. Identification of Four Polyhydroxyalkanoate Structural Genes in Synechocystis Cf. Salina PCC6909: In Silico Evidences. J Proteomics Bioinform 2016, 09 (02), 28–37. 10.4172/jpb.1000386.

(17) Guzman, L. M.; Belin, D.; Carson, M. J.; Beckwith, J. Tight Regulation, Modulation, and High-Level Expression by Vectors Containing the Arabinose P(BAD) Promoter. J Bacteriol 1995, 177 (14), 4121–4130. 10.1128/jb.177.14.4121-4130.1995.

(18) Huang, C.; Wang, J.; Liu, H.; Huang, R.; Yan, X.; Song, M.; Tan, G.; Zhi, F. Ketone Body β-Hydroxybutyrate Ameliorates Colitis by Promoting M2 Macrophage Polarization through the STAT6-Dependent Signaling Pathway. BMC Med 2022, 20 (1), 148. 10.1186/s12916-022-02352-x.

(19) Feng, G.; Wu, Z.; Yang, L.; Wang, K.; Wang, H. β-Hydroxybutyrate and Ischemic Stroke: Roles and Mechanisms. Mol Brain 2024, 17 (1), 48. 10.1186/s13041-024-01119-0.

(20) Goldberg, E. L.; Shchukina, I.; Asher, J. L.; Sidorov, S.; Artyomov, M. N.; Dixit, V. D. Ketogenesis Activates Metabolically Protective Γδ T Cells in Visceral Adipose Tissue. Nat Metab 2020, 2 (1), 50–61. 10.1038/s42255-019-0160-6.

(21) Shichita, T.; Sugiyama, Y.; Ooboshi, H.; Sugimori, H.; Nakagawa, R.; Takada, I.; Iwaki, T.; Okada, Y.; Iida, M.; Cua, D. J.; Iwakura, Y.; Yoshimura, A. Pivotal Role of Cerebral Interleukin-17–Producing ΓδT Cells in the Delayed Phase of Ischemic Brain Injury. Nat Med 2009, 15 (8), 946–950. 10.1038/nm.1999.

(22) Benakis, C.; Brea, D.; Caballero, S.; Faraco, G.; Moore, J.; Murphy, M.; Sita, G.; Racchumi, G.; Ling, L.; Pamer, E. G.; Iadecola, C.; Anrather, J. Commensal Microbiota Affects Ischemic Stroke Outcome by Regulating Intestinal Γδ T Cells. Nat Med 2016, 22 (5), 516–523. 10.1038/nm.4068.

(23) Stefan, V. E.; Weber, D. D.; Lang, R.; Kofler, B. Overcoming Immunosuppression in Cancer: How Ketogenic Diets Boost Immune Checkpoint Blockade. Cancer Immunol Immunother 2024, 74 (1), 23. 10.1007/s00262-024-03867-3.

